# Combination of Imipridone ONC201 or ONC206 with Temozolomide and Radiotherapy in triple ITR therapy reduces intracranial tumor burden and prolongs survival in an orthotopic wild-type IDH GBM mouse model

**DOI:** 10.1101/2024.09.25.610187

**Authors:** Lanlan Zhou, Leiqing Zhang, Jun Zhang, Laura Jinxuan Wu, Shengliang Zhang, Andrew George, Marina Hahn, Howard P. Safran, Clark C. Chen, Attila A. Seyhan, Eric T. Wong, Wafik S. El-Deiry

**Affiliations:** Laboratory of Translational Oncology and Experimental Cancer Therapeutics, Warren Alpert Medical School, Brown University, Providence, RI, United States; Department of Pathology and Laboratory Medicine, Warren Alpert Medical School, Brown University, Providence, RI, United States; Joint Program in Cancer Biology, Lifespan Health System and Brown University, Providence, RI, United States; Legorreta Cancer Center at Brown University, Providence, RI, United States; Hematology/Oncology Division, Department of Medicine, Lifespan Health System and Brown University, Providence, RI, United States; Department of Neurosurgery, Lifespan and Brown University, Providence, RI; Departments of Medicine, Radiation Oncology, Neurosurgery and Neurology, Brown University, Providence, RI

**Keywords:** Glioblastoma multiforme, IDH wild-type, imipridones, ONC201, ONC206, MGMT, Temozolomide, Radiotherapy, integrated stress response, TRAIL

## Abstract

Glioblastoma remains the most lethal common primary brain tumor in adults with limited therapeutic options. TIC10/ONC201, a first-in-class imipridone we discovered, achieved meaningful therapeutic effects in phase I/II trials in patients with diffuse gliomas harboring H3K27M mutations, and currently the drug is in randomized phase III testing (ACTION trial; NCT05580562). ONC201 targets mitochondrial protease ClpP to disrupt oxidative phosphorylation and trigger the integrated stress response (ISR), TRAIL/DR5, and tumor cell death. We hypothesized that ONC201 and its analogue ONC206 synergize with temozolomide (TMZ) and ionizing radiation (IR), standard-of-care glioblastoma therapies. ONC201 enhances TMZ or IR-induced apoptosis, and cytotoxicity. ClpP-silencing suppresses ONC201-induced cytotoxicity but not TMZ or RT. Both ONC201 and ONC206 reduce expression of TMZ-resistance mediator MGMT. Suppression of MGMT protein was observed in H3K27M-mutated DIPG cell lines following treatment with ONC201 or ONC206 with or without TMZ. Cytokine profiling indicates distinct ONC201 alterations relative to TMZ suggesting distinct anti-tumor immune mechanisms. Triple IR+TMZ+ONC201 (ITR) therapy prolongs median survival to 123 days with a tail on survival curve (3-of-7 mice alive beyond 200-days) in an orthotopic U251 GBM model versus ONC201 (44-days; p=0.000197), IR (63-days; p=0.0012), TMZ (78-days; p=0.0354), ONC201+IR (55-days; p=0.0004), ONC201+TMZ (80-days; p=0.0041) and IR+TMZ (103-days; p>0.05). By 231-days, the only surviving mice were in IRT group. Our results support investigation of ONC201/ONC206 in combination with TMZ and IR (ITR) in GBM or H3K27M mutated diffuse glioma therapy.

## Introduction

CNS tumors are among the most lethal cancers in children and adults. There are more than 13,000 new cases of malignant brain tumors diagnosed each year in the US alone. Glioblastoma (GBM) is the most common, fast growing, and aggressive primary malignant tumor in the CNS in adults [1, 2] carrying a poor prognosis, with a median survival of less than 15 months with only 10% of patients responding to standard of care [3]. GBM is a fast-growing and aggressive brain tumor that accounts for ∼60 – 70% of malignant gliomas and ∼15% of CNS tumors [4]. Moreover, diffuse intrinsic pontine glioma (DIPG), another lethal pediatric brainstem tumor primarily associated with the H3K27M mutation, displays a dire 5-year survival rate of less than 1%, primarily associated with the H3K27M mutation [5]. The standard of care (SOC) for GBM is maximally safe surgical resection, followed by concurrent radiation therapy (RT) and temozolomide (TMZ) for 6 weeks, then adjuvant TMZ for 6 months, yet the overall survival (OS) rates remain relatively low. Radiation is the primary treatment modality for DIPG but the benefit from concurrent temozolomide is questionable in light of higher systemic toxicities [6], while chemotherapies are offered to infants upfront to delay radiation until their brains become more developed.

TMZ, an alkylating agent, exerts its therapeutic action by methylating DNA adenine and guanine residues on single-stranded DNA [7]. The O-6-methyl-guanine (O-6-MG) when methylated, if not repaired, can be mis-paired to induce base transition mutations [8]. If single- or double-stranded DNA breaks occur and are unrepaired, this ultimately results in growth arrest or apoptosis [9]. The presence or absence of O6-methylguanine-DNA methyltransferase (MGMT), an enzyme involved in DNA repair, diminishes or enhances TMZ’s efficacy in tumors with unmethylated and methylated promoter of this enzyme [7]. GBM with methylated MGMT that inhibits MGMT protein expression has significantly longer OS compared to those that are unmethylated when treated with radiation and temozolomide [10].

ONC201 (also known as TIC10, originally discovered in the El-Deiry Lab [11, 12] as a TRAIL-inducing compound), is a novel first-in-class small molecule imipridone compound, that has shown promise in targeting various pathways involved in cancer progression, such as the extrinsic apoptotic pathway, upregulation of pro-apoptotic TRAIL receptor DR5 [13–15]. ONC201 was originally found to inactivate ERK/AKT leading to Foxo3a nuclear translocation and TRAIL gene activation [11]. Later it was discovered that ONC201 activates the integrated stress response (ISR) leading to ATF4-dependent upregulation of TRAIL receptor DR5 [16]. Other studies have documented depletion of cancer stem cells [17] and activation of the immune response mediated by natural killer cells that, in part, involve TRAIL/DR5 as a component of the host innate immune response against cancer [15]. Recent investigations have revealed that ONC201, along with its imipridone analogs ONC206 and ONC212, bind to mitochondrial ClpP as an agonist [18, 19], inhibiting oxidative phosphorylation as a component of its anti-cancer mechanism, eventually leading to apoptosis [12, 20]. Moreover, ONC201 and its analogs exhibit effectiveness against DIPG cell lines and cross the blood-brain-tumor barrier, suggesting their clinical applicability against CNS tumors.

Clinical studies and compassionate use cases of imipridone ONC201 have highlighted its promising efficacy, particularly in H3K27M-mutated tumors, including DIPG and GBM. Current trials exploring ONC201 as well as its analogs in combination with standard therapies seek to enhance their therapeutic potential in these aggressive tumors. Importantly, ONC201 is currently being explored in clinical studies in patients with various solid tumors and a randomized phase III study in patients with recurrent H3K27M-mutated diffuse glioma (ACTION trial; NCT05580562). ONC206 is currently being investigated in two Phase 1 studies in children and young adults with newly diagnosed or recurrent diffuse midline glioma (NCT04541082, NCT04732065). Notably, in recent compassionate use cases for ONC201, patients with H3K27M mutations in different brain tumors, particularly DIPG, glioblastomas, and other high-grade midline brain tumors have shown promising results, including significant tumor shrinkage and clinical improvements [21]. Furthermore, ONC201 and its analogs trigger ISR and this could attenuate the expression of DNA-damage repair proteins [22], which could potentiate the effects of radiation and concomitant TMZ for MGMT unmethylated GBM. The clinical experience with ONC201 in H3K27M–mutant pediatric DIPG provides further rationale for treatment with TMZ in combination with RT to investigate synergy in GBM and DIPG cells, *in vivo* models, and patients.

We investigated the synergistic effects of imipridone ONC201 in combination with RT and TMZ (ITR) as a triple combination therapy against GBM. Through *in vitro* and *in vivo* experiments on various brain tumor cell lines, including GBM, DIPG, and atypical teratoid rhabdoid tumors, we assessed cell viability, ISR activity, and apoptosis. Findings from *in vivo* orthotopic GBM mouse studies demonstrated that the triple combination treatment significantly prolongs survival and reduces tumor burden, decreases cell proliferation, and induces more apoptosis compared to single or dual therapies, indicating a promising avenue for GBM frontline treatment. We extended our studies and show similar effects with ONC206 in ITR combination therapy. Importantly we observed unique cytokine profiles and suppression of MGMT expression in glioma cell lines by imipridones ONC201 and ONC206 as potential synergy mechanisms with TMZ. Suppression of MGMT protein was observed in H3K27M-mutated DIPG cell lines following treatment with ONC201 or ONC206 with or without TMZ. Overall, our preclinical findings support further exploration of the ONC201 and ONC206 ITR regimen as a potential treatment for GBM and diffuse gliomas with H3K27M mutations.

## Results

### The combination of imipridone ONC201 and RT produces synergy with loss of cell viability in GBM cell lines

Previously, single-agent cytotoxicity of imipridone ONC201 or in combination with RT or TMZ was demonstrated *in vitro* [22–24]. We therefore hypothesized that ONC201 may synergize with both RT and TMZ in therapy of malignant brain tumors. To test this hypothesis, glioblastoma (GBM: SNB19, T98G and U251), diffuse intrinsic pontine glioma (DIPG: SF8628), and atypical teratoid rhabdoid tumor (ATRT: BT-12, BT-16) cell lines were tested using cell viability or colony formation assays with ONC201 up to 20 μM alone or in combination with radiotherapy up to 8 Gy or temozolomide up to 100 μM [24]. We used Western blot analysis to document drug-induced apoptosis of treated cells with single or combination therapy which showed synergy between ONC201 and RT and between ONC201 and TMZ with the best combination indices of 0.51 and 0.21 respectively [24].

We later investigated whether the combination of ONC201 and RT produced synergistic effects similar to what was reported previously [24]. To investigate the synergistic activity of ONC201 in combination with RT and against solid tumor cell lines, we treated a panel of human brain tumor cell lines including SNB19 (GBM), U251 (GBM), and atypical teratoid rhabdoid tumor (ATRT: BT-12, BT-16) cell lines with varying doses of TRAIL pathway inducer ONC201 and RT and evaluated the cell viability and colony formation after treatment. Data from cell viability demonstrated synergy between ONC201 and RT in all cell lines tested (**Fig. 1A)**. As shown in **Fig. 1A**, the cell lines with the highest synergy scores were GBM cell lines SNB19, and U251 cells as well as the ATRT BT16 cell line.

**Figure 1.**
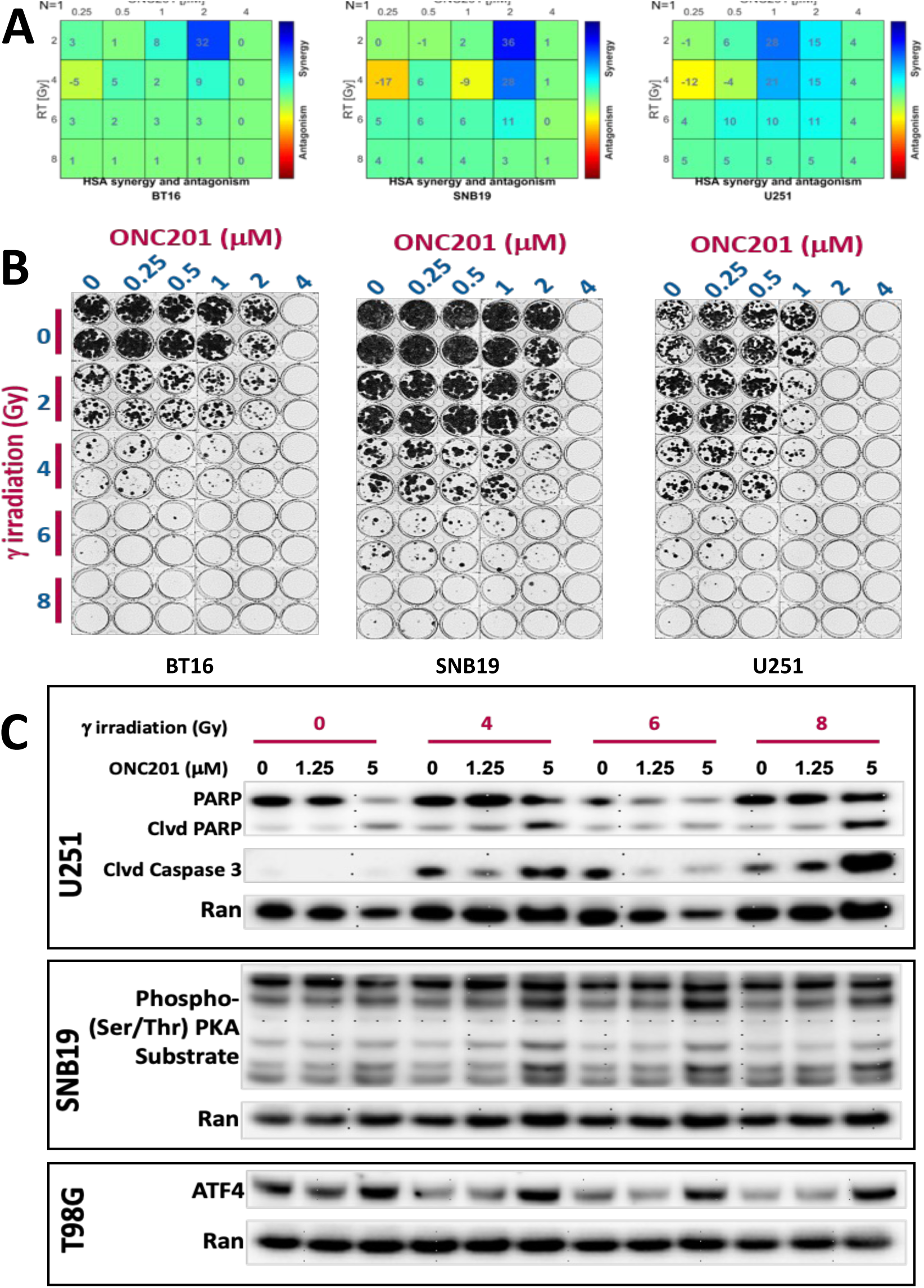
ONC201 synergizes with RT. **A.** Combination of ONC201 and RT with GBM cell lines SNB19, T98G, and U251, diffuse intrinsic pontine glioma (DIPG) cell line SF8628 and atypical teratoid rhabdoid tumor (ATRT: BT-12, BT-16) cell lines show treatment effects on cell viability, **B.** Colony formation assays with ONC201 up to 4 μM alone or in combination with radiotherapy up to 8 Gy. Colony formation and CellTiter Glo assays were performed to determine the cell viability after treatment. Highlighted in boxes are dose combinations that show synergy. Data demonstrate synergy between ONC201 and radiation therapy. **C.** Western blots were utilized to assess induction of PKA substrate phosphorylation, induction of ATF4 as a marker of ISR activation, and multiple markers of cell death such as PARP and cleaved PARP, cleaved Caspase 3 in treated brain tumor cells.

### The combination of ONC201 and RT synergistically suppresses colony formation by GBM cell lines

After observing synergistic cytotoxicity across GBM cell lines in short-term cell viability assays, we explored whether similar sensitivity and synergy could be observed under longer-term exposure. A 7-day colony formation assay was conducted to assess the effects of ONC201 alone (up to 4 μM) or in combination with RT (up to 8 Gy) (**Fig. 1B**). The LD90 was found under the exposure of ONC201 and radiation at 2 μM and 4 Gy for BT16, 2 μM and 6 Gy for SNB19, and 1 μM and 4 Gy for U251, respectively (**Fig. 1B**). BT16 was most sensitive followed by U251 and then SNB19 for reduction in colony-forming ability with combined ONC201 and RT treatment as compared to single agent treatment (**Fig. 1B**). Thus, the long-term colony formation assay in **Fig. 1B** demonstrates potent anti-cancer effects when ONC201 is combined with RT for the treatment of GBM cells.

### ONC201 and RT activate ISR, enhance PARP and caspase cleavage and apoptosis

Having observed high levels of synergy across all tested GBM cell lines, we explored whether the treatment with ONC201 and RT induced apoptosis. When T98G, SNB19 and U251 glioma cell lines were exposed to various doses of radiation, Western blot analysis showed an ONC201 dose-dependent increase in the expression and activation levels of ATF4 protein, a mediator of ISR and a marker of ISR activation, possibly by specific eIF2 alpha kinases as described earlier [16], Increased PKA substrate phosphorylation was also observed at the highest ONC201 dose level of 5 μM, and markers of apoptosis including PARP and cleaved caspase 3 in treated GBM cell lines SNB19, T98G and U251 (**Fig. 1C**). Overall, the data presented in **Fig. 1C** supports the conclusion of synergy between ONC201 and radiation therapy in brain tumors.

### Imipridone ONC201 synergizes with Temozolomide against GBM and DIPG cell ines

Having observed single-agent cytotoxicity with monotherapy with ONC201 and TMZ [24], we investigated further whether the combination of ONC201 and TMZ produced synergistic effects. We performed both cell viability and colony formation assays to evaluate the combination of ONC201 and TMZ with the diffuse intrinsic pontine glioma (DIPG) cell line SF8628, and GBM cell line U251 cell lines using increasing concentrations of both drugs in combination. As shown in **Fig. 2A** and **B** a CI <1 was noted across all concentrations of ONC201 and TMZ tested, suggesting some degree of synergy in these brain tumor cell lines using cell viability and colony formation assays.

**Figure. 2.**
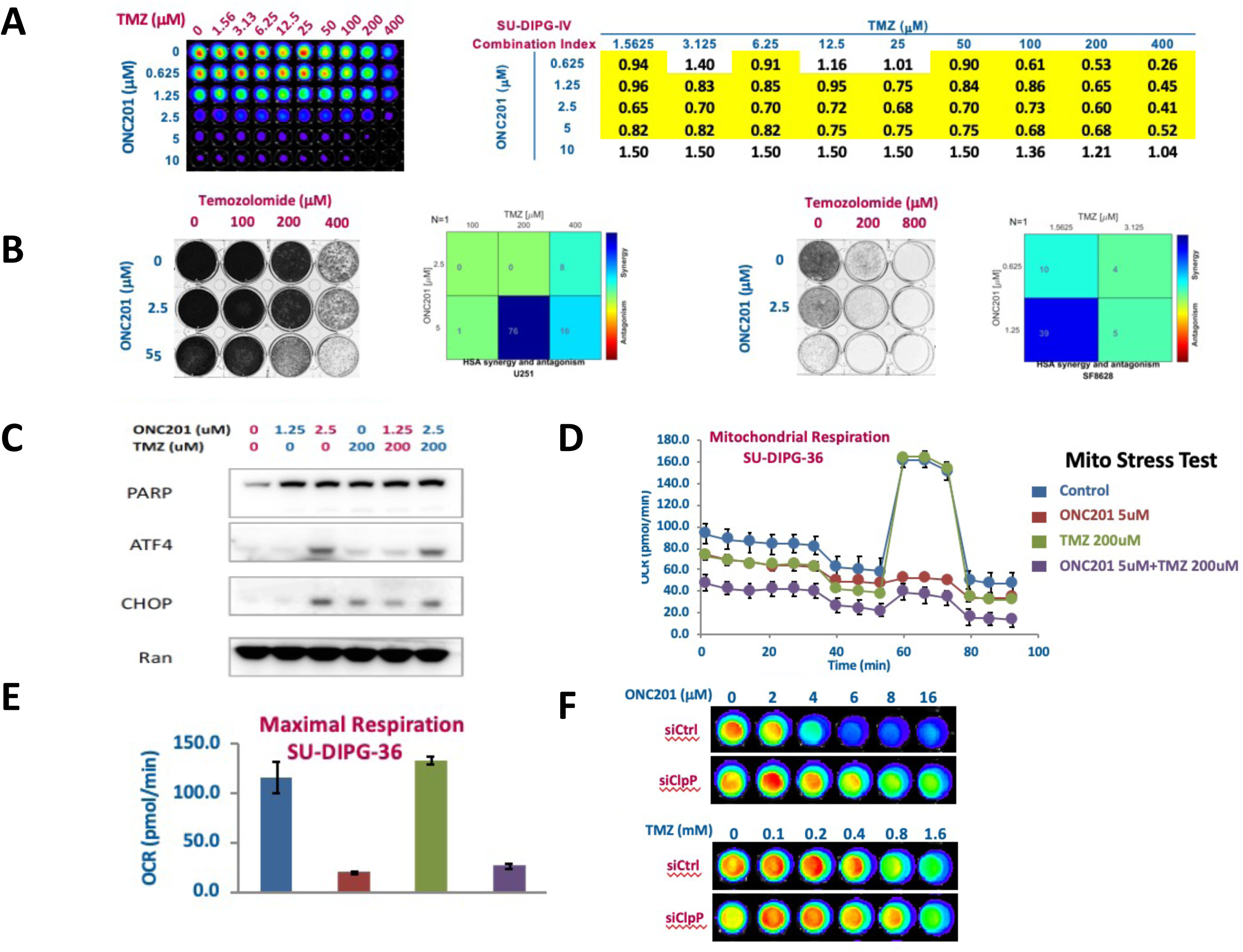
ONC201 synergizes with TMZ. **A**. The combination of ONC201 and TMZ with the diffuse intrinsic pontine glioma (DIPG) cell line SF8628 and GBM cell line U251 using cell viability. CellTiter Glo assays were performed to determine the cell viability after treatment. Combination indices (CI) were calculated by the method of Chou and Talalay using the CompuSyn software. Highlighted in boxes are dose combinations that show synergy. Data demonstrate synergy between ONC201 and TMZ up to 400 μM. **B.** Colony formation assays were performed to determine long-term cell viability after treatment. Highlighted in boxes are dose combinations that show synergy. Data demonstrate synergy between ONC201and TMZ therapy up to 400 μM. **C.** Western blotting was utilized to assess induction of ATF4 and CHOP as markers of ISR activation, and cleaved PARP as a marker of cell death in treated brain tumor cells. Data demonstrate synergy between ONC201 and TMZ up to 200 μM. **D.** Mitochondrial dysfunction induced by ONC201 is reflected in reduction of maximal cell respiration in the SU-DIPG 36 cell line. Treatment conditions shown in different colors as indicated. **E.** Effect of ONC201 on oxygen consumption rate (OCR) tested in SU-DIPG 36 cell line using single-agent therapy with ONC201 or TMZ or doublet therapy using ONC201 in combination with TMZ. OCR of treated cells was measured by Seahorse Analyzer. Treatment conditions as in panel D. **F.** siRNA knockdown of ClpP in GBM cell line shows that ONC201 induced cytotoxicity depends on ClpP whereas siRNA knockdown of ClpP has no effect on TMZ-induced cytotoxicity.

### Imipridone ONC201 in combination with TMZ induces ISR and apoptosis in U251 GBM cells

After observing significant levels of cytotoxic synergy in GBM cell lines, we investigated the potential activation of the ISR and induction of apoptosis following treatment with ONC201 and TMZ. Western blot analysis showed that PARP was upregulated upon treatment with either ONC201 or TMZ, but ATF4 was induced only at the highest ONC201 concentration of 2.5 μM (**Fig. 2C**). However, CHOP, a marker of ISR activation and inducer of the apoptotic pathway, could be induced at a lower concentration of ONC201 in combination with TMZ (**Fig. 2C**). Together, these findings demonstrate a robust synergy when ONC201 was combined with TMZ, leading to PARP cleavage during apoptosis involving upregulation of ATF4 and CHOP from ISR activation.

### ClpP activation by imipridone ONC201 impairs oxidative phosphorylation and induces apoptosis following reduction of respiratory chain complex subunits

Based on findings of ISR activation and induction of apoptosis following treatment with ONC201 and TMZ, we then evaluated the relevance of mitochondrial protease ClpP (caseinolytic mitochondrial matrix peptidase proteolytic subunit that degrades damaged or misfolded proteins activation), a recently described binding and activation target of ONC201 [18, 19], on oxidative phosphorylation and mitochondrial function in DIPG, GBM and ATRT cell lines following treatment with ONC201 and TMZ. Our data demonstrate that treatment with ONC201 or ONC201 in combination with TMZ results in ClpP activation, decreased basal oxygen consumption rate and spare reserve capacity in the SU-DIPG 36 cell line and that mitochondrial dysfunction induced by ONC201 is reflected in reduction of maximal cell respiration in these cell lines (**Fig. 2D)**. Notably, TMZ alone has no effect on the basal oxygen consumption rate in the DIPG cell line (**Fig. 2D, E)**.

### Imipridone ONC201-induced cytotoxicity depends on ClpP whereas TMZ-induced cytotoxicity is independent of ClpP

To further decipher the mechanism behind the decreased basal oxygen consumption rate and spared respiratory reserve capacity from ONC201 with or without TMZ, we investigated the activation of mitochondrial protease ClpP in the SU-DIPG 36 cell line. RNA interference studies were conducted using control versus ClpP siRNAs that were introduced into SU-DIPG-36 cells at concentrations of ONC201 and TMZ ranging from 0 to 16 μM and 0 to 1.6 mM, respectively. As expected, siClpP protected cells from loss of viability after ONC201 but not with TMZ treatment (**Fig. 2F**). We observed that ClpP activation increases the production of mitochondrial reactive oxygen species (ROS) in DIPG cells treated with control siRNA whereas DIPG cells treated with ClpP specific siRNA were strongly protected from ONC201-mediated cytotoxicity suggesting that the activity of ONC201 depends on ClpP (**Fig. 2F)**. The result confirms that mitochondrial protease ClpP is an activation target of ONC201 and indicates that ONC201-induced DIPG cell death is probably mediated by the production of mitochondrial reactive oxygen species (ROS) in DIPG cells. By contrast, temozolomide induced cytotoxicity does not depend on ClpP.

### ONC201 synergizes with both RT and TMZ as ITR combination therapy in the rhabdoid tumor cell line BT16

We hypothesized that ONC201 may synergize TMZ or RT in brain tumors. To test this hypothesis, we evaluated the combination of ONC201, RT and TMZ (named as ONC201 ITR therapy) with rhabdoid tumor cell line BT16 using cell viability, colony formation, and Western blot assays. CellTiter Glo and colony formation and assays were performed to determine cell viability and synergy after treatment of the BT16 cell line with the ITR combination of ONC201, TMZ, and RT and found that triple combination exerts strong synergy as indicated by inhibition of cell viability (**Fig. 3A**) and suppression of colony formation (**Fig. 3B**). Quantification of dose combinations showed strong synergy between ONC201, TMZ, and RT (**Fig. 3C**). Collectively, ONC201 synergizes with TMZ and RT in cell viability (**Fig. 3A**) and colony formation assays (**Fig. 3B**) in rhabdoid tumor cell line BT16.

**Figure 3.**
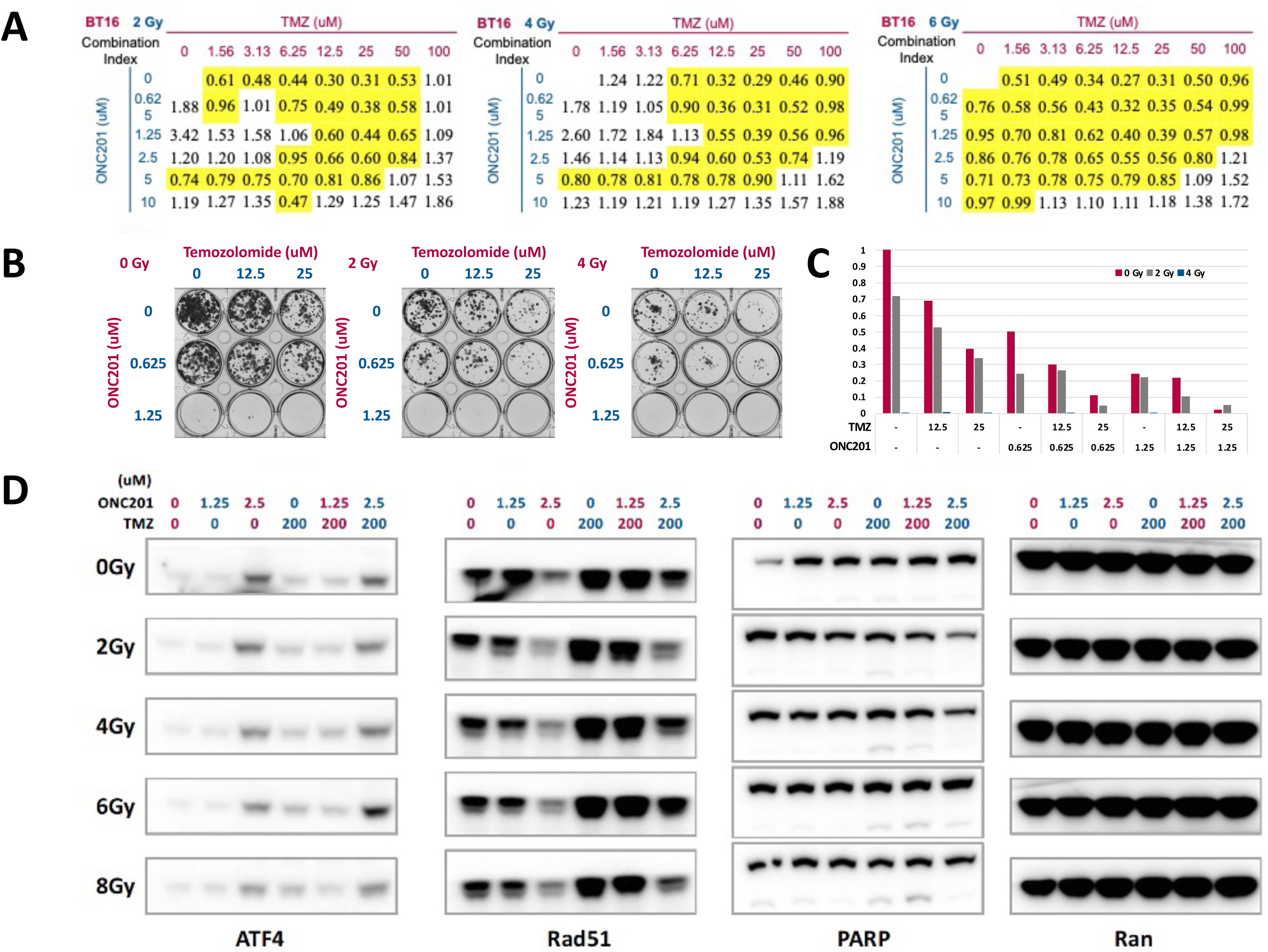
ITR triple combination of ONC201 with RT and TMZ synergizes in rare rhabdoid tumor BT16 cells. ITR triple combination of ONC201, RT, and TMZ with BT16 cells was assessed using cell viability, colony formation, and Western blotting. A. Combination indices are shown for rhabdoid tumor cell line BT16 treated at different radiation doses, TMZ and ONC201 as indicated. B. Colony formation assays with ONC201 alone or in combination with TMZ at indicated concentrations of each drug are shown. C. Quantification of colony formation of GBM cells after ITR triple combination treatment shows triple drug synergy. D. Western blotting to assess induction of ATF4 and CHOP as markers of ISR activation, cleaved PARP as marker of cell death and suppression of Rad51, a selective DNA repair target to radio-sensitize glioma stem cells in treated brain tumor cells. Data demonstrate strong synergy with ITR therapy with ONC201 RT, and TMZ.

Western blots analysis was utilized to assess the apoptosis of treated cells and showed induction of ATF4 as marker of ISR activation, cleaved PARP as marker of cell death, and suppression of ATPase Rad51, a selective DNA repair target that radio-sensitizes glioma stem cells in treated brain tumor cells [25] (**Fig. 3D**). Rad51 inhibition removes SOX2-expressing cells and abolishes clonogenicity. Rhabdoid tumor cell line BT16 shows an increase in Rad51 expression after RT and TMZ treatment and Rad51 levels are inversely correlated with radiosensitivity. Downregulation of Rad51 after ONC201 exposure markedly increases the cytotoxicity of TMZ. These data demonstrate a strong synergy between ONC201 and TMZ plus RT (**Fig. 3C**), and this synergy is a result of induction of ISR, and subsequent apoptosis.

### ITR triple combination of ONC201, TMZ, and RT shows strong synergy in BT16, SNB19, and U251 GBM cell lines, induction of both ISR and apoptosis

We sought to explore whether the synergy from ITR triple therapy that resulted in the induction of ISR and apoptosis is a generalizable phenomenon in other GBM cell lines. We therefore applied the combination of ONC201, TMZ and RT to SNB19, T98G, U138, and U251 glioblastoma cells and performed Western blot analysis of the expression of PARP, ATF4, LC3B and ClpX (**Fig. 4**). Indeed, we found that ONC201 ITR triple combination induced ATF4 and inhibited ClpX, suggesting that ISR was activated and ClpP was most likely unleashed in all GBM cell lines tested. In addition, microtubule-associated protein light chain 3 (LC3), which serves as a specific marker for autophagy, had no significant changes with treatments. Collectively, our biomarker studies demonstrated that ITR triple combination treatment decreases tumor cell proliferation, induces more apoptosis, and inhibits ClpX to unleash mitochondrial ClpP (**Fig. 4**).

**Figure 4.**
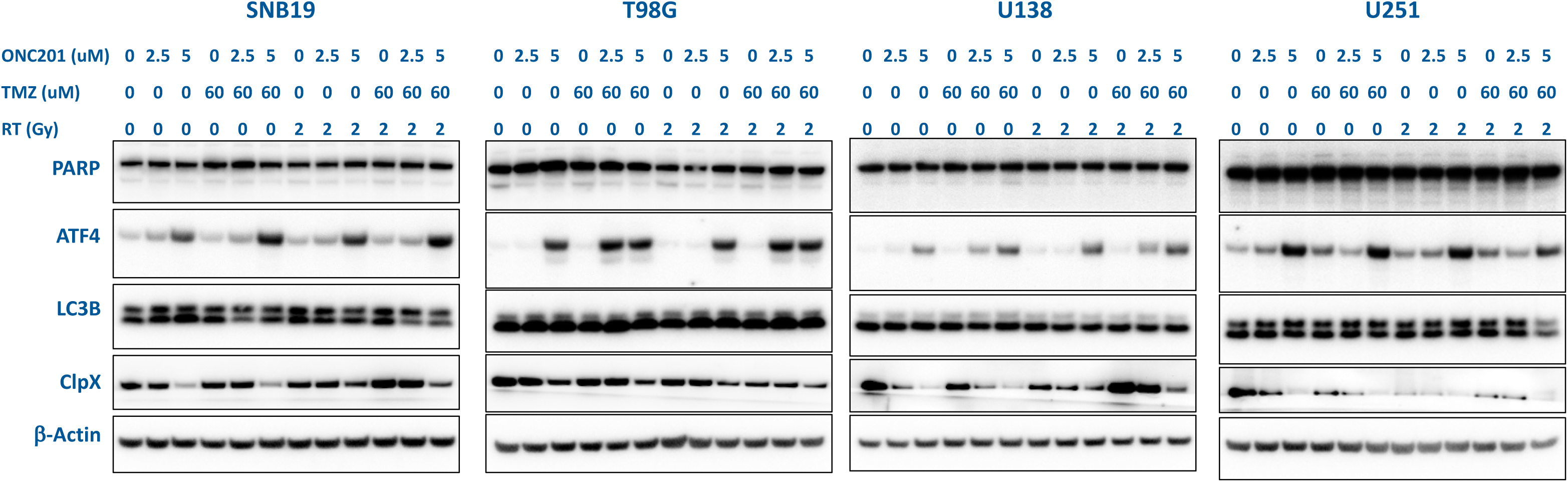
ITR triple combination of ONC201, RT and TMZ shows strong synergy, induces ATF as marker of ISR activation, and inhibits ClpX to unleash ClpP in SNB19, T98G, U138, and U251 GBM cells. ITR combination treatment effects with ONC201, RT, and TMZ on BT16, SNB19, and U251 GBM cell lines as assessed by Western blotting. ITR combination treatment induces ATF4 and inhibited ClpX. Microtubule-associated protein light chain 3 (LC3) used as a specific marker to monitor autophagy shows no significant changes with treatments. ClpX and LC3 is correlative of synergy. Cells treated with ONC201, TMZ, and RT at indicated drug concentrations or radiation doses at 37°C for 48 hours.

### ONC201 or ONC206 ITR therapy shows synergies with RT and TMZ associated with distinct cytokine profiling

We treated GBM tumor cells (U251) with 10 μM ONC201 or 1 μM ONC206 as ITR therapy in combination with 200 μM TMZ and 2 or 10 Gy of radiation for 48 h and subsequently analyzed the cell culture supernatant using Luminex 200 technology (**Fig. 5**). Several cytokines, chemokines, and growth factors associated with angiogenesis and/or EMT were found to be downregulated in the U251 cell line with both ONC201 and ONC206. Notably, angiopoietin-1 (a potent angiogenic growth factor that is critical for vessel maturation, adhesion, migration, and survival), M-CSF (a cytokine that regulates the proliferation, differentiation, and functional activation of monocytes), beta 2-microglobulin, PDL-1, IFN-g, FAS, CCL2, CCL13, all had decreased secretion post-treatment with ONC201 in combination with TMZ and RT at both 2 and 10 Gy radiation doses. A similar but less pronounced decrease in these cytokines, chemokines, and growth factors was seen with ITR therapy using ONC206 in combination with TMZ and RT at both 2 and 10 Gy radiation doses. Likewise, several cytokines, chemokines, and growth factors associated with immunosuppression, including angiopoietin-1, Fas, and soluble PD-L1, were also downregulated post-treatment. Furthermore, ONC206 but not ONC201 in combination with TMZ and RT induced the secretion of TRAIL R3, CD40/TNFRSF5, CCL3/MIP-1a, IL-18/IL-1F4, GD-15, CXCL5/ENA-78, IL-8/CXCL8, IL2, VEGF, angiopoietin-2 (an angiogenic growth factor that promotes cell death and disrupts vascularization), and TRAIL R2.

**Figure 5.**
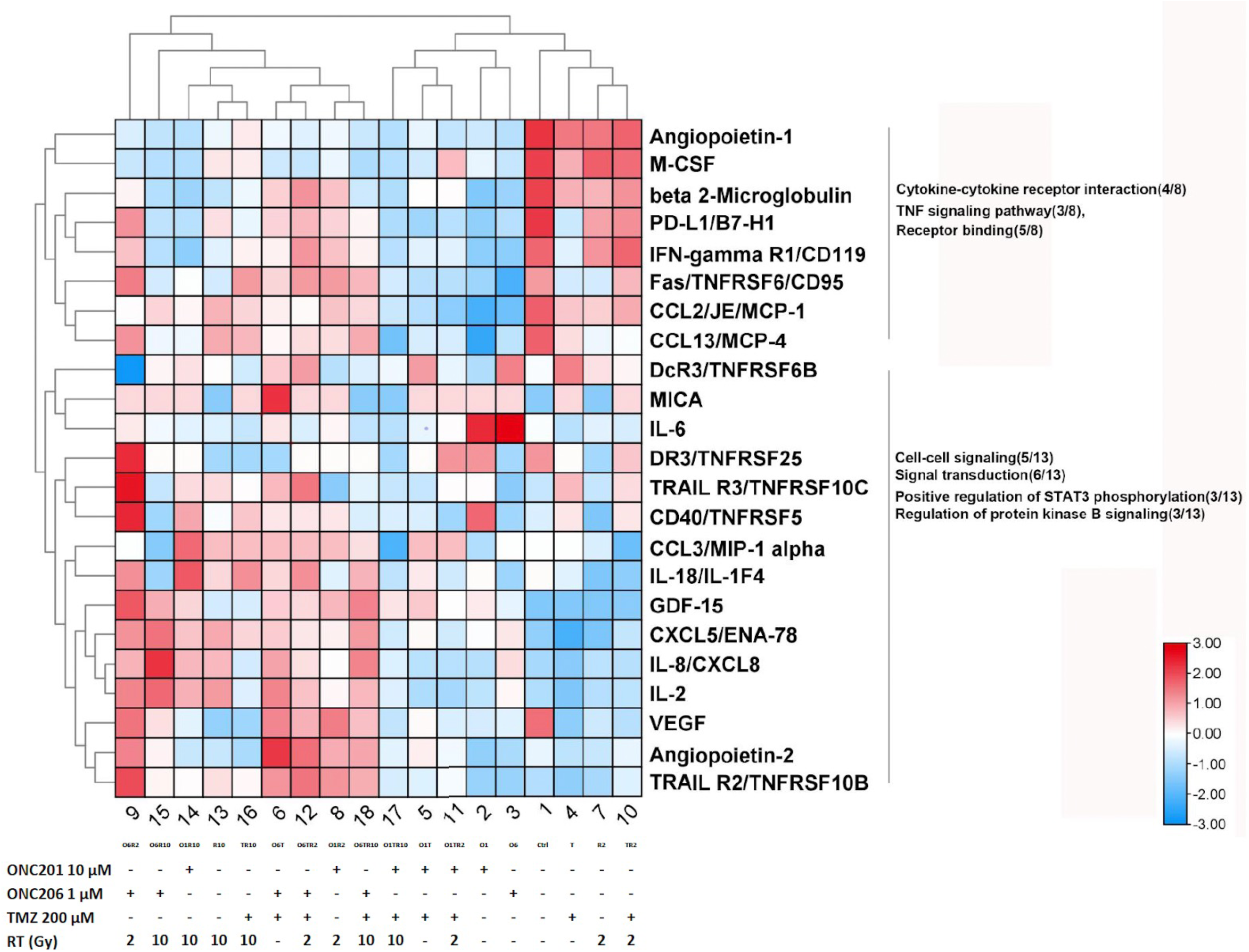
Differential cytokine profiling of ITR triple combinations of ONC201 or ONC206 with RT and TMZ for 4hr in U251 GBM cell line *in vitro*. U251 tumor cells treated with ONC201 or ONC206, TMZ, and RT at indicated concentrations or radiation doses for 48 h and cell culture supernatant analyzed with the Luminex 200. Fold-change is shown where red indicates a positive fold-change and green indicates a negative fold-change (N = 3).

### ONC201 ITR in combination with TMZ and RT is an effective antitumor agent in mice with orthotopic GBM

Our cell culture studies above showed that ITR therapy with ONC201 synergizes with TMZ and RT in GBM cell lines SNB19, T98G, U138, and U251 and rhabdoid tumor cell line BT16. ITR-treated cells induce ATF4 and CHOP as markers of ISR activation, cleave PARP or caspase 3 as markers of apoptosis following ITR triple treatment regimen in cell culture experiments. We next sought to explore whether this ITR triple treatment also shows a strong synergy and induction of ISR and apoptosis in an orthotopic GBM mouse model following treatment with ONC201 in combination with TMZ and RT.

Using intracranial injection of luciferase-expressing U251 GBM cells with a KOPF model 940 small animal stereotaxic frame and a Stoelting Quintessential Stereotaxic Injector, we confirmed tumor formation and growth using bioluminescence imaging. Mice were randomized to receive weekly treatment of ONC201 (100 mg/kg p.o.) and/or radiotherapy (2 Gy local irradiation) and/or TMZ (20 mg/kg i.p.) for two weeks for short-term biomarker studies or four weeks for long-term survival and tumor monitoring (**Table 1**, **Fig. 6A)**. In longer-term *in vivo* experiments, Kaplan Maier survival curve shows that, a four-week ITR triple therapy treatment with ONC201 in combination with TMZ and RT increases median survival of mice to 123 days as compared to double-agent or single-agent treatment regimens in an aggressive intracranial xenograft of human GBM U251-Luc cells **(Fig. 6A)**. Effectively, ONC201 cooperated with TMZ and RT to triple the survival duration of such brain tumor-bearing mice and reduce tumor size as compared to the control group **(Fig. 6A** and **B).**

**Figure 6.**
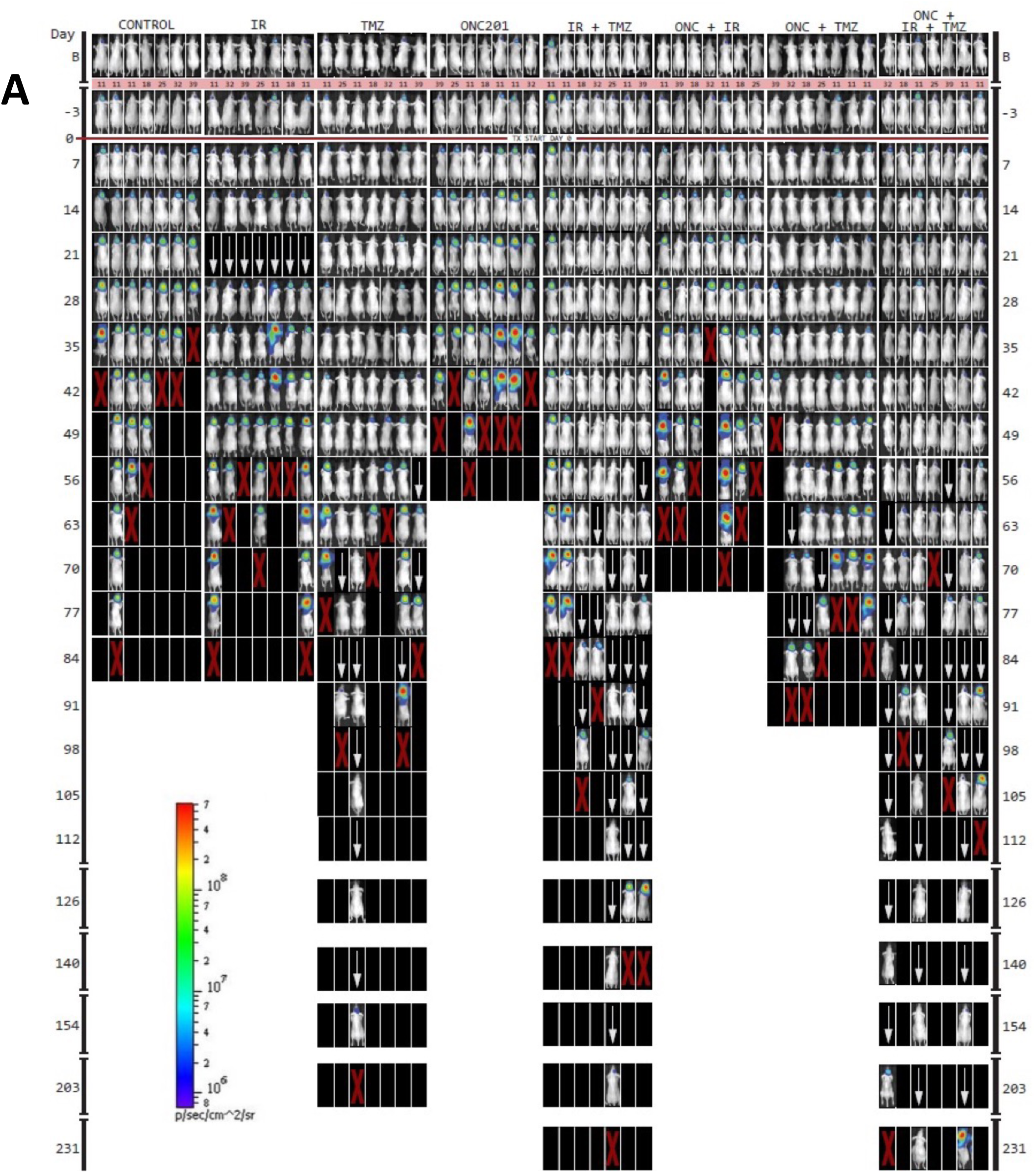
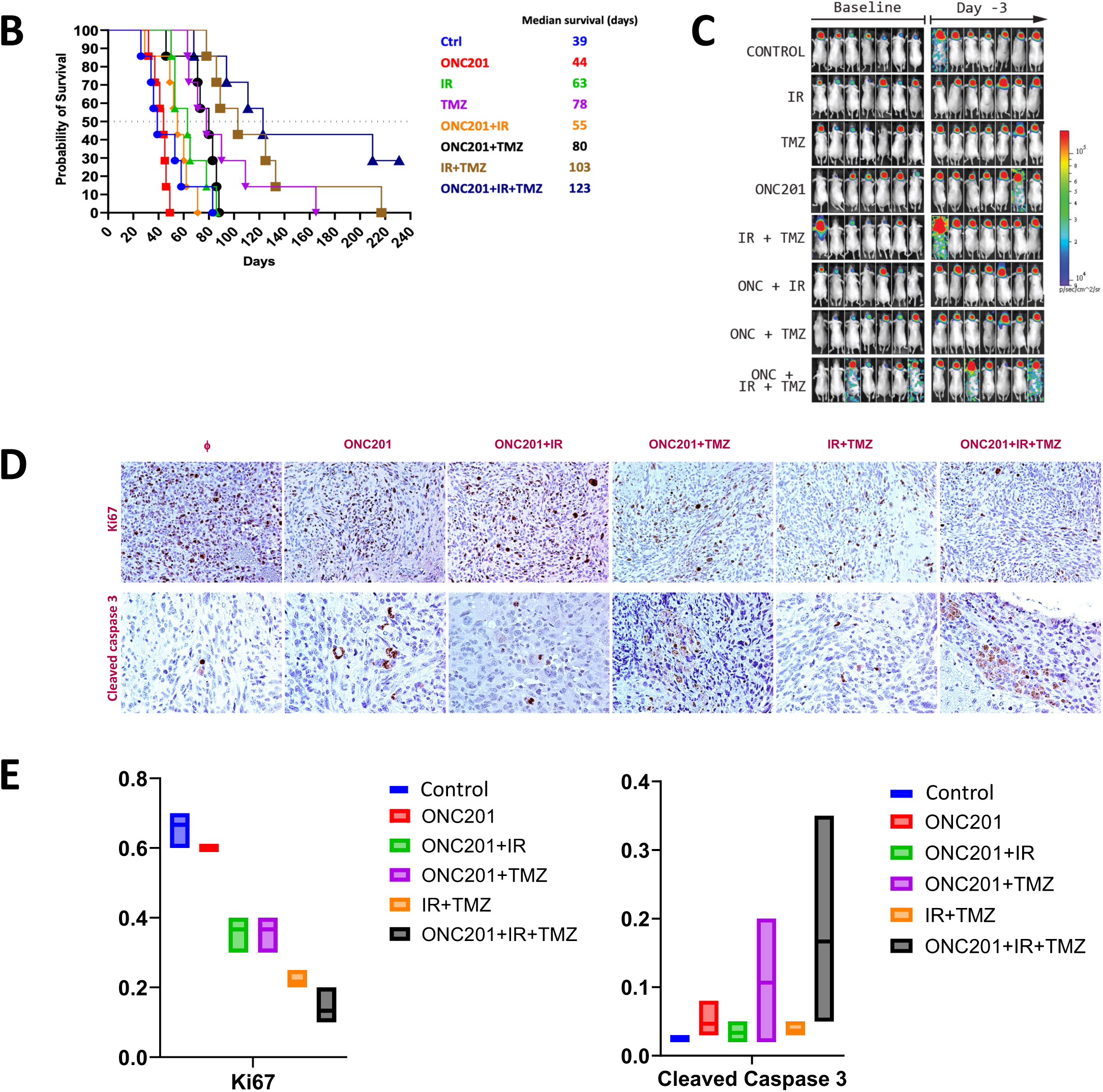
ITR triple combination of ONC201, radiotherapy, and TMZ for 4 weeks significantly prolongs mouse survival and reduces tumor burden in an orthotopic brain tumor model. **A.** Randomized treatment group mice received weekly treatment of ONC201 (100 mg/kg p.o.) and/or radiotherapy (2 Gy local irradiation) and/or TMZ (20 mg/kg i.p.) for four weeks for long-term survival and tumor monitoring. **B.** Long-term survival studies show that ITR triple combination of ONC201, RT, and TMZ significantly prolongs survival and reduces tumor burden as compared to single- and dual-treatments. **C.** Tumors were growing at the time just before treatment was initiated: tumor images at baseline and day −3. Long-term survival studies show tumors were growing at the time just before treatment was initiated. **D.** IHC assessment of ITR triple combination of ONC201, radiotherapy, and TMZ for 2 weeks shows reduced Ki67 expression and induction of cleaved caspase 3. **E.** Quantification of short-term biomarkers demonstrating ITR triple combination treatment decreases tumor cell proliferation (Ki67) and induces more apoptosis (cleaved caspase 3).

**Table 1.** Design of short and long-term ITR treatment *in vivo* studies.

| Table 1. Design of *in vivo* studies |  |  |  |
| --- | --- | --- | --- |
| Groups | 4-week treatments for survival<br>2-week treatments for biomarkers |  |  |
|  | ONC201<br>100 mg/kg weekly,<br>p.o. | TMZ<br>20 mg/kg weekly,<br>i.p. | IR<br>2Gy weekly, local |
| Control | - | - | - |
| ONC201 | + | - | - |
| TMZ | - | + | - |
| IR | - | - | + |
| ONC201+TMZ | + | + | - |
| TMZ+IR | - | + | + |
| ONC201+IR | + | - | + |
| ONC201+TMZ+IR | + | + | + |

Tumor formation and growth were confirmed with bioluminescence imaging. Mice were randomized to receive weekly treatment of ONC201 (100 mg/kg p.o.) and/or radiotherapy (2 Gy local irradiation) and/or TMZ (20 mg/kg i.p.) for two weeks for short-term biomarker studies or four weeks for long term survival and tumor monitoring as previously described and illustrated in **Table 1**. Findings from this study confirmed that tumors were growing at the time just before ONC201 ITR treatment was initiated by tumor images at baseline and day −3 (**Fig. 6C**). Thus, our longer-term survival studies show that the ITR triple combination of ONC201, radiotherapy, and TMZ significantly prolongs survival and reduces tumor burden as compared to single-treatment and dual-treatment combinations. Triple therapy with ITR prolonged median survival to 123 days with 3 of 7 mice alive beyond 200 days in an orthotopic U251 glioblastoma model relative to ONC201 (44 days; p=0.000197), IR (63 days; p=0.0012), TMZ (78 days; p=0.0354), ONC201 + IR (55 days; p=0.0004). ONC201 + TMZ (80 days; p=0.0041) and IR + TMZ (103 days; p>0.05) (**Supplementary Figure 1**). By 231 days, the only surviving mice were in the IRT group. These results support further larger *in vivo* studies, consideration for ONC201 or ONC206 into the current standard of care for glioblastoma in the upfront setting, and also suggest potential synergies of triple therapy in tumors with H3K27M mutations that should be further explored in preclinical and clinical studies.

### An ITR triple combination of ONC201, RT, and TMZ for 2 weeks inhibits proliferation and induces apoptosis in an orthotopic brain tumor model

In addition to the long-term survival studies, we sought to explore a short-term cell proliferation (Ki67) and apoptosis markers in tumor specimens collected from mice two weeks after the start of a single, double, or ITR triple therapy with ONC201, TMZ, and/or RT. Short-term biomarker studies show that treatment with ITR triple combination of ONC201, RT, and TMZ for two weeks inhibited tumor cell proliferation and induced apoptosis assessed by measurements of Ki67 and cleaved caspase 3 respectively by immunohistochemical analysis of tumor specimens collected two weeks after the start of treatment (**Fig. 6D and E**). Thus, these biomarker studies demonstrate that ITR triple combination treatment decreases tumor cell proliferation and induces more apoptosis compared to single-treatment or dual-treatment combinations.

### ITR triple combination therapy with ONC206, RT, and TMZ gives similar *in vivo* reduction in proliferation and induction of apoptosis as compared to ONC201 ITR therapy

Given that ONC206 has entered clinical trials for patients with brain tumors (NCT04732065, NCT04541082) we sought to gather preliminary data on its potential to synergize with RT and TMZ *in vivo* as we demonstrated above with ONC201. Mice that were orthotopically implanted with brain tumor cells (as in **Figure 6**) received weekly treatment of ONC206 (100 mg/kg p.o.) and/or radiotherapy (2 Gy local irradiation) and/or TMZ (20 mg/kg i.p. weekly) for two weeks for short-term biomarker studies. The short-term biomarker studies (**Figure 7**) demonstrate that ITR triple combination of ONC206, radiotherapy, and TMZ for two weeks decreases tumor cell proliferation (Ki67) and induces apoptosis (cleaved Caspase 3). These results were similar to what we observed with the triple combination of ONC201, RT, and TMZ suggesting that either of the two imipridones ONC201 or ONC206 could be investigated further in the clinic in combination with RT and TMZ.

**Figure 7.**
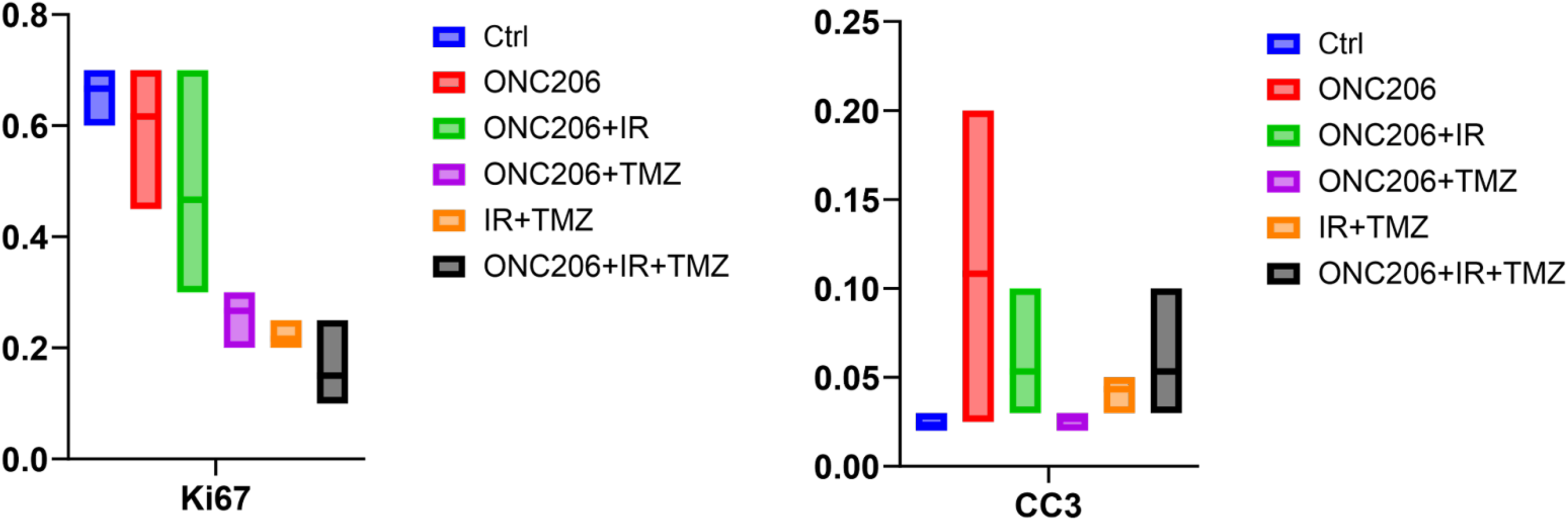
ITR triple combination of ONC206 with RT and TMZ for two weeks inhibits proliferation and induces apoptosis in the orthotopic U251 GBM model *in vivo*. *A g*roup of three mice received weekly treatment of ONC206 (100 mg/kg p.o. weekly) and/or radiotherapy (2 Gy local irradiation) and/or TMZ (20 mg/kg i.p. weekly) for two weeks for short-term biomarker studies. Short-term biomarker studies demonstrate that ITR triple combination of ONC206, radiotherapy, and TMZ for two weeks decreases tumor cell proliferation (Ki67) and induces apoptosis (cleaved Caspase 3). Quantification is shown from three mice per group.

### Reduction of expression of TMZ-resistance mediator MGMT in H3K27M-mutated DIPG cell lines following treatment with ONC201 or ONC206 with or without TMZ

We recently reported that expression of MGMT in our commonly used GBM cell lines is low as compared to other solid tumor cell lines where it is quite high and that TMZ can have effects on tumor cell killing by T-cells within a few hours before there is time for cell division or mutations to occur [26]. MGMT methylation leading to loss of MGMT protein expression is common in GBM and associated with response to TMZ [27, 28]. By contrast, MGMT is rarely if ever methylated in H3K27M-mutated diffuse gliomas leading to TMZ resistance [29, 30]. We hypothesized that effects of imipridones on the ISR and global inhibition of protein translation might impact on MGMT expression and potential synergies when combined with TMZ for treatment of glioma cells. Because our GBM cell lines including the U251 cells used in our orthotopic brain tumor experiments have low or undetectable MGMT expression, we chose to test effects of imipridones on MGMT expression when used alone or in combination with TMZ in treatment of DIPG cells (**Fig. 8**). We found evidence that imipridones (ONC201, ONC206 and ONC212) modulate MGMT and ClpX expression in DIPG cell lines. Both ONC201 and ONC206 reduced MGMT expression in SF8628 cells either in the absence or presence of TMZ (**Fig. 8A**). We investigated the effects of the imipridones on ClpX and MGMT in additional DIPG cell lines including SU-DIPG-13, SU-DIPG-25, SU-DIPG-36, and SU-DIPG-IV (**Fig. 8B-F**). We found that to some extent there was reduction of MGMT protein expression in the DIPG cell lines treated by the imipridones alone or in combination with TMZ. These novel findings suggest a potential mechanism of synergy between imipridones and TMZ through suppression of MGMT protein expression in DIPG cells that are typically unmethylated for MGMT and resistant to effects of TMZ.

**Figure 8.**
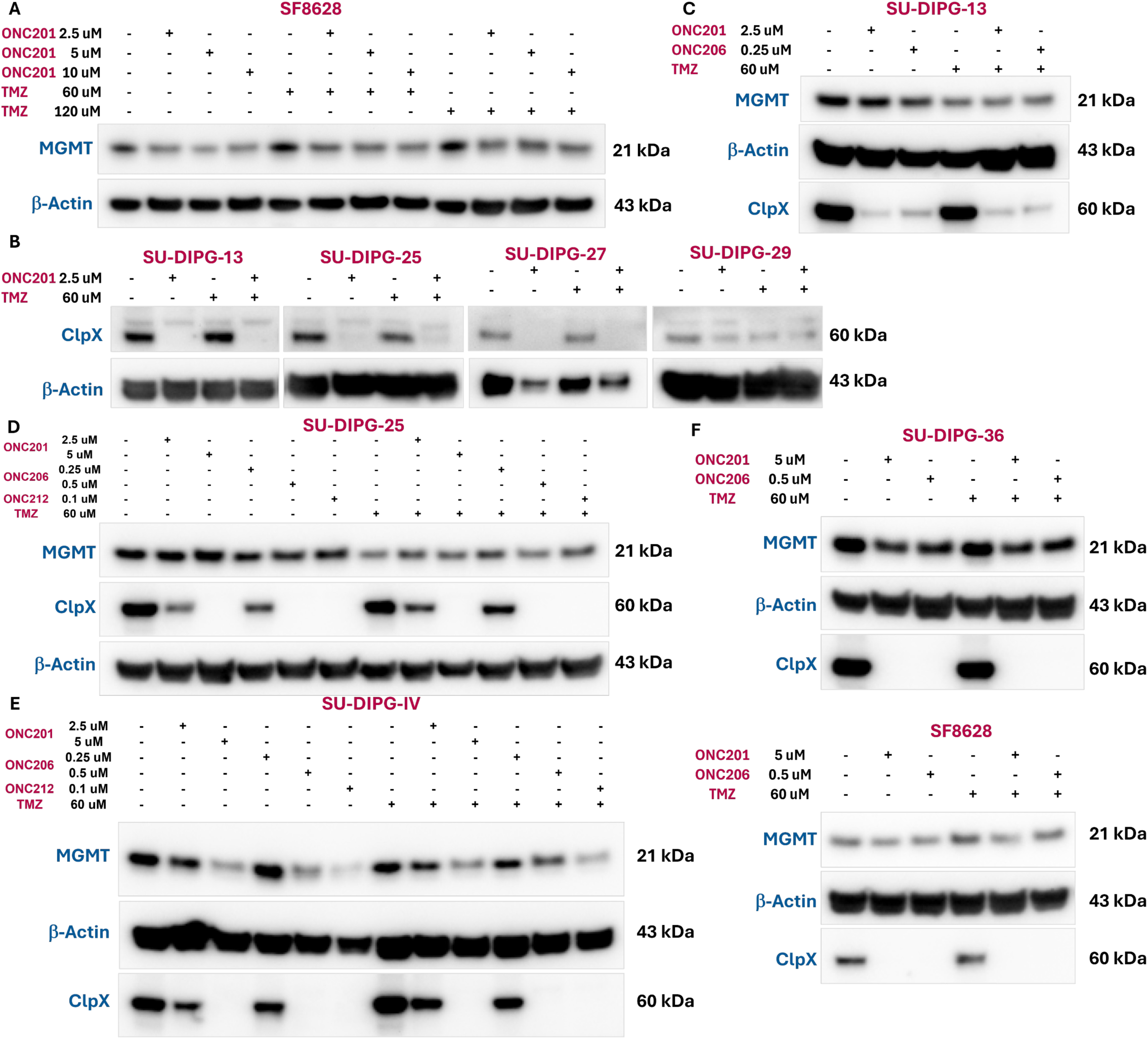
Imipridones (ONC201, ONC206 and ONC212) modulate MGMT and ClpX expression in DIPG cell lines. **(A)** ONC201 reduced MGMT expression in SF8628 cell line. **(B)** ONC201 inhibited ClpX expression in multiple DIPG cell lines (SU-DIPG-13, SU-DIPG-25, SU-DIPG-27 and SU-DIPG-29). Imipridones inhibited ClpX expression in a dose-dependent manner but did not reduce MGMT in SU-DIPG-13 **(C)** and SU-DIPG-25 **(D)** cell lines. **(E)** Imipridones decrease ClpX and MGMT expression in a dose-dependent manner in the SU-DIPG-IV cell line. **(F)** Imipridones (ONC201 and ONC206) inhibit ClpX expression and reduce MGMT in SU-DIPG-36 and SF8628 cell lines.

## Discussion

Despite the utilization of multimodal therapies including RT and chemotherapy involving TMZ, the prognosis of GBM remains dismal, with consolidated treatment yielding a median survival of 14.6 months [31]. Furthermore, several randomized clinical trials investigating agents targeting EGFR, mTOR, and VEGF, along with multiple vaccine studies and immune checkpoint inhibitors, have yielded disappointing results after promising early-stage (phase I/II) studies in GBM. Our *in vitro and in vivo* findings demonstrate for the first time that ITR therapy combining ONC201 or ONC206 with TMZ and RT may be a safe and effective therapeutic strategy for GBM. Similar triple therapy using a putative TRAIL agonist TRA-8 in combination with TMZ and RT also resulted in enhanced cytotoxicity against glioma stem cells [32]. Their kinome analysis revealed that this enhanced cytotoxicity appears to be mediated by inhibition of JAK-STAT and Src tyrosine kinase signaling. WE have identified a novel mechanism for synergy between imipridones and TMZ involving suppression of MGMT expression that likely occurs through the imipridone-induced ISR with suppression of global protein translation. Our results and those of Arafat *et al*. [32] indicate that TRAIL is an important tumoricidal effector for glioblastoma. However, our results differ from those from Arafat *et al*. [32] by showing for the first time the effect on MGMT expression after imipridone treatment through the unique mechanism of imipridones that differs from TRAIL receptor agonists.

Our experimental findings from ITR triple combination treatment of GBM cell lines demonstrate that ITR triple combination treatment of ONC201 or its analog ONC206 in combination with TMZ and RT significantly inhibits cell survival, induces more apoptosis, disrupts mitochondrial function, inhibits various cytokines, chemokines, and growth factors associated with angiogenesis and/or EMT, immunosuppression or promotes factors associated with cell death and disrupts vascularization. Moreover, our *in vivo* studies using the orthotopic U251 GBM mouse model randomized to receive weekly ITR treatment of ONC201 or ONC206 and/or radiotherapy and/or TMZ for four weeks for either long-term survival and tumor monitoring or two weeks for short-term biomarker studies demonstrate that the ITR triple combination treatment significantly prolongs survival and reduces tumor burden compared to single and dual treatment combinations in the long-term studies. The ITR triple combination not only reduces tumor cell proliferation, but also induces more apoptosis, and inhibits ClpX to unleash mitochondrial ClpP in short-term biomarker studies. The effect of imipridones on MGMT in multiple DIPG cell lines provides a mechanism by which their combination with TMZ could overcome TMZ resistance in the clinic. This should be further tested across tumor types including H3K27M-mutated gliomas.

Our studies demonstrate that ONC201 synergizes with RT, induces apoptosis as indicated by the generation of multiple markers of cell death such as PARP, cleaved PARP, and cleaved Caspase 3, induces PKA substrate phosphorylation, upregulates ATF4 as a marker of ISR activation in treated brain tumor cells.

We further investigated the relevance of mitochondrial protease ClpP, a recently described binding target of ONC201 [18, 19], on oxidative phosphorylation and mitochondrial function in DIPG cells. Our findings reveal that mitochondrial dysfunction induced by ONC201 is reflected in reduction of maximal cell respiration in brain tumor cells. The observed mitochondrial protein degradation and cytotoxicity attributed to ONC201 is dependent on ClpP as knockdown of ClpP by siRNA protects multiple human cancer cell lines from ONC201-mediated cytotoxicity. Notably, temozolomide alone does not have an effect on the basal oxygen consumption rate. We demonstrated that ITR triple treatment of BT16 tumor cells with ONC201, TMZ and RT shows induction of ATF4 indicative of ISR activation, cleaved PARP as a marker of apoptosis, and suppression of RAD51, a selective DNA repair target to radio-sensitize glioma stem cells in treated brain tumors. We observed that ITR triple combination treatment induces PKA substrate phosphorylation, induces ATF4/ISR activation, and inhibits ClpX to unleash ClpP, and multiple markers of cell death in treated in SNB19, T98G, U138, and U251 GBM cell lines.

Cytokine profiling from ITR-treated U251 GBM cells with ONC201 or ONC206 in combination with TMZ and RT for 48 hr revealed that several cytokines, chemokines, and growth factors associated with angiogenesis and/or EMT were downregulated with both ONC201 and ONC206. Likewise, several cytokines, chemokines, and growth factors associated with immunosuppression were downregulated post-treatment, including angiopoietin-1, Fas, and soluble PD-L1. Interestingly, ONC206 but not ONC201 in combination with TMZ and RT induced the secretion of TRAIL R3, CD40/TNFRSF5, CCL3/MIP-1a, IL-18/IL-1F4, GD-15, CXCL5/ENA-78, IL-8/CXCL8, IL2, VEGF, angiopoietin-2, and TRAIL R2.

Long-term survival studies showed that ITR triple combination of ONC201, radiotherapy, and TMZ significantly prolongs survival and reduces tumor burden as compared to single-treatment and dual treatment combinations. We carefully documented that tumors were growing at the time just before ITR or other treatments were initiated by capturing tumor images at baseline and day −3. Evaluation of tumor images of surviving mice treated with indicated agents or a combination of agents on days 130 and 161 after implantation demonstrate that ONC201 synergizes with TMZ and RT and prolongs survival better that single or double treatments in the human GBM U251 orthotopic mouse model. Biomarker studies demonstrate that ITR triple combination of ONC201, RT, and TMZ for two weeks inhibits proliferation and induces apoptosis in tumor specimens. The biomarker studies demonstrate that ITR triple combination treatment also inhibits ClpX to unleash mitochondrial ClpP. The short-term *in vivo* ITR treatment study with ONC206 in combination with TMZ and RT achieves similar anti-tumor effects as evaluated by assessment of markers of cell proliferation (Ki67) and apoptosis (cleaved caspase 3) in tumor specimens collected two weeks after the start of treatment in the orthotopic U251 GBM model *in vivo*.

Based on our preclinical cell culture and *in vivo* data, ITR triple therapy with ONC201 or ONC206 synergizes with the TMZ and RT in an orthotopic glioma mouse model serves as a step towards clinical translation and the studies provide pharmacodynamic biomarkers.

Our findings demonstrate for the first time that ITR therapy combining ONC201 or ONC206 with TMZ and RT may be a safe and effective therapeutic strategy for GBM. Future studies may unravel novel signaling mechanisms following ONC201 or ONC206 plus TMZ and RT treatment of humanized and syngeneic immunocompetent *in vivo* model of GBM, and GBM patients. This includes the role of the ISR, TRAIL pathway, mitochondrial metabolism, ClpP, and MGMT as well as potential immune effects in the efficacy of the combination therapy. Furthermore, findings from this study might establish the ITR combination regimen of ONC201 or ONC206 in combination with TMZ and RT in tumors with or without H3K27M mutation and MGMT gene methylation [33] for targeted therapy.

Limitations include observed synergistic effects may be specific to certain cell lines, raising concerns about generalizability across different GBM tumor types including GBM tumor cells, organoids, and *in vivo* models with and without the H3K27M mutation and MGMT gene methylation. Mechanistic insights, while partially provided by Western blot analysis, require further studies such as transcriptomics or RNA-Seq to fully elucidate molecular pathways and consider potential off-target effects. Additional limitations include lack of use of syngeneic orthotopic mouse models and patient-derived xenograft humanized immunocompetent orthotopic GBM models. Lack of immune cell subset analysis, such as T-cell and NK-cell populations, and patient-derived models, highlights the need for validation across a broader spectrum of GBM subtypes and immune cell subtypes. The complexity of TME, including extracellular matrix, stromal cells, immune cells, fibroblasts, cytokines, tumor-associated myeloid cells/macrophages, T lymphocytes, B lymphocytes, Natural Killer cells, Neutrophils, Dendritic cells, GBM cells/Glioma stem cells among others [34] may significantly influence treatment responses.

While promising, additional research is necessary to confirm and translate these findings into clinical practice. Our data suggest synergy between the ITR triple therapy (i.e. imipridone ONC201 or ONC206, radiotherapy, and temozolomide), including a tail on the survival curve not seen with mono- or dual-therapies. These findings support further exploration of mechanisms, PK, PD biomarkers, activity *in vivo*, and translation to the clinic.

Of note we found synergy between ONC201 and TMZ in H3K27M-mutated SF8628 cells and have previously reported TMZ effects on immune killing by T-cells of high MGMT-expressing tumor cells [26]. We showed that in SU-DIPG-36 cells that ONC201 suppresses oxygen consumption rate observed with TMZ alone, likely due to suppression of oxidative phosphorylation by ONC201. We also discovered the effect of imipridones on suppressing MGMT expression to potentially sensitize tumor cells with unmethylated MGMT to the anti-tumor effects of TMZ. Thus, the synergies and potentially beneficial therapeutic effects of combining ONC201 or other imipridones with TMZ, in addition to their combined use in the ITR combination could be further exploited. Our results support further studies combining ONC201 or ONC206 with TMZ and RT (ITR) in glioblastoma as well as H3K27M mutated diffuse gliomas.

## Materials and Methods

### Cell culture and reagents

Human brain tumor cell lines used in this study include GBM cell lines SNB19, T98G, and U251, diffuse intrinsic pontine glioma (DIPG) cell line SF8628, and atypical teratoid rhabdoid tumor (ATRT) cell lines BT-12 and BT-16. These cell lines were obtained from the American Type Culture Collection (ATCC). SU-DIPG 36 cells were a gift from Michelle Monje (Stanford University). All cell lines were confirmed to be free of mycoplasma contamination using PCR testing methods. All cell lines were cultured in their ATCC-recommended media supplemented with 10% fetal bovine serum (FBS) and 1% penicillin/streptomycin at 37°C in a 95% humidified atmosphere containing 5% carbon dioxide. Chemotherapy agents used in the study were ONC201 (obtained form Chimerix), solubilized in DMSO at a storage concentration of 20 mM; and TMZ was purchased from Selleckchem (Catalog No. S1237), reconstituted in DMSO at a storage concentration of 20 mM.

### Measurement of cell viability

Cells were seeded at a density of 3 × 10^3^ cells per well in a 96-well plate (Greiner Bio-One, Monroe, NC, USA). Cell viability was assessed using the CellTiter Glo assay (Promega, Madison, WI, USA). Cells were mixed with 25 μL of CellTiter-Glo reagents in 100 μL of culture volume, and bioluminescence imaging was measured using the Xenogen IVIS imager (Caliper Life Sciences, Waltham, MA). The percent of cell viability was calculated by normalizing the luminescence signal to control wells and was reported as % viability ± SD.

Dose-response curves were generated, and the half-maximal inhibitory concentration (IC-50) was calculated using GraphPad Prism version 9.2.0 (RRID: SCR_002798). For IC50 determination, concentrations were log-transformed, and data were normalized to the control, followed by a log (inhibitor) versus response (three parameters) analysis as described previously [35].

The combination indices (CI) were calculated using CompuSyn software (ComboSyn, Inc.). CI values below 1 indicate some degree of synergy, while values below 0.5 indicate very strong synergy as described previously [35].

### Collection of cell supernatant samples for cytokine profiling

Cells were plated at 3.5 × 10^4^ cells in a 48-well plate (Thermo Fisher Scientific, Waltham, MA, USA) in complete medium and incubated at 37°C with 5% CO2. At 24 hours after plating, almost all the tumor cells were adherent to the bottom of the flask and the complete medium was removed and replaced with the drugged medium. After 48 hours of treatment, cell culture supernatants were collected and were frozen at −80°C until the measurement of cytokines, chemokines, and growth factors was performed. On the day of analysis, samples were thawed and centrifuged to remove cellular debris.

### Cytokine profiling of drug-treated tumor cells

An R&D systems Human Premixed Multi-Analyte Kit (R&D Systems, Inc., Minneapolis, MN) was run on a Luminex 200 Instrument (LX200-XPON-RUO, Luminex Corporation, Austin, TX) per the manufacturer’s guidelines.

Sample levels of a custom panel of analytes (**Fig. 5**) including Angiopoietin-1, Angiopoietin-2, M-CSF, beta-2-Microtubulin, PDL1/B7-H1, IFN-gamma, FAS, CCL2, CCL3, CCL13, DrC3, MICA, IL-6, CD40, IL18, GDF-15, CXCL5, IL-8, IL2, VEGF, TRAIL-R2, and TRAIL-R3 were measured for the drug treatment experiments. Analyte values were reported in picograms per milliliter (pg/mL).

### Knockdown of ClpP gene expression by siRNA transient transfection

Cells were seeded in a 96-well plate and transfected transiently with control and target-specific siRNAs using lipofectamine RNAi MAX (Invitrogen) following the manufacturer’s instructions and cells were cultured for 24 hours. After 24 hours, cell culture media was replaced with fresh media and treated with ONC201 or TMZ at indicated concentrations and further cultured for an additional 48 hours.

### Western blot analysis

Cells were lysed in RIPA buffer (Sigma-Aldrich, St. Louis, MO, USA) containing cocktail protease inhibitors (Roche, Basel, Switzerland). Equal amounts of cell lysates were electrophoresed through 4-12% SDS-PAGE and then transferred to PVDF membranes. The transferred PVDF membranes were blocked with 5% skim milk at room temperature, then incubated with primary antibodies indicated in a blocking buffer at 4°C overnight. Antibody binding was detected on PVDF with appropriate HRP-conjugated secondary antibodies by a Syngene imaging system (Syngene, Bangalore, India).

### Colony formation assays

Approximately 300 brain tumor cells per well were cultured in 12-well plates. Cells were treated with different drugs or with radiation therapy at various doses for three days, then the cells were cultured with a drug-free complete medium for two weeks with the fresh medium being changed every three days. The cell colonies were fixed with 10% formalin and stained with 0.05% crystal violet at the end of the experiments.

### Knockdown of gene expression by siRNA transient transfection

Cells were seeded in a 96-well plate and transfected with siRNA using lipofectamine RNAi MAX (Invitrogen) as described in the protocol. 24 hours later after siRNA transfection, cells were co-cultured with cancer cells carrying luciferase reporter for three days.

### Luciferase reporter assay

Brain tumor cells carrying a constitutive luciferase (Luc) reporter were treated with different drugs or with radiation therapy at various doses and the luciferase reporter expression in cancer cells was examined based on bioluminescence using the IVIS imaging system (PerkinElmer, Hopkin, MA, USA).

### *In vivo* experiments

The experimental protocol for the in vivo study (Protocol # 22-02-0004) was approved by the Institutional Animal Care and Use Committee of Brown University (Providence, RI, USA).

### Intracranial injection

With a KOPF model 940 small animal stereotaxic frame and a Stoelting Quintessential Stereotaxic Injector, NCRNU-Female, 5 weeks of age athymic nude mice (Taconics, https://www.taconic.com/mouse-model/ncr-nude) were used to inject 2×10^5^ U251-Luc expressing cells in 5 mL PBS orthotopically into the frontal region of the cerebral cortex a site 2.5 mm lateral (right), 1.5 mm anterior, and 2.5 mm ventral with respect to the bregma to produce a GBM tumor. ONC201 was prepared at 25 mg/mL concentration in 10% DMSO + 20% Cremophor EL + 70% PBS. TMZ was prepared. at 2 mg/mL in 5% DMSO + 30% PEG300 + 65% ddH2O.

### Treatment regimen

Cohorts of mice (n =7) were treated with vehicle control, single agent with ONC201 (100 mg/kg weekly, p.o.), TMZ (20 mg/kg 5x/week, i.p.), RT (IR) (2Gy 5x/week, local), as well as with dual therapy with ONC201 + TMZ, and a triple therapy with ONC201 + TMZ + RT. All mice were imaged using the IVIS® Spectrum in vivo imaging system for bioluminescence after tumor cell injection to monitor tumor growth. All mice were followed daily to record survival. At the end of the 4 weeks or longer treatment period depending on the tumor burden, the effect of treatments on survival was quantified by calculating the time until drug-treated and control mice develop terminal symptoms and are euthanized. For survival studies, morbidity criteria were used. Survival data were plotted on a Kaplan-Meier curve.

### Survival analysis

Overall survival of orthotopic xenograft mice harboring U251-Luc intracranial tumors treated with a single dose of vehicle, ONC201 (100 mg/kg weekly, p.o), TMZ (20 mg/kg 5x/week, i.p.), RT (IR) (2Gy 5x/week, local), or with dual therapy with ONC201 + TMZ, and triple therapy with ONC201 + TMZ + RT at indicated time points after implantation was analyzed by plotting the survival data on a Kaplan Maier survival curve.

### Tumor size assessment

Representative images of survived mice treated with the indicated drug on days 130 and 161 after implantation. The relative tumor size measured by bioluminescence imaging using the IVIS® Spectrum in vivo imaging system in an orthotopic xenograft of U251-Luc in mice treated with a single dose of vehicle, ONC201, TMZ, RT (IR), as well as with dual therapy with ONC201 + TMZ, and a triple therapy with ONC201 + TMZ + RT. The bioluminescence scale bar on the right applies to all images in the panel. Data demonstrate that ONC201 synergizes with TMZ and RT in the human GBM U251 orthotopic mouse model.

### Immunohistochemistry

Excised tumor tissue samples were fixed using 10% neutral buffered formalin and embedded in paraffin. Tissue sections of approximately 5 µm were cut with a microtome and mounted on glass microscope slides for subsequent staining.

Hematoxylin and eosin (H&E) staining was performed for all tumor tissue specimens. Paraffin embedding and sectioning of slides were performed by the Brown University Molecular Pathology Core Facility. Slides were dewaxed in xylene and subsequently hydrated in ethanol at decreasing concentrations. Antigen retrieval was carried out by boiling the slides in 2.1 g citric acid (pH 6) for 10 min. Endogenous peroxidases were quenched by incubating the slides in 3% hydrogen peroxide for 5 min. After nuclear membrane permeabilization with Tris-buffered saline plus 0.1% Tween 20, slides were blocked with horse serum (Cat# MP-7401-15, Vector Laboratories, Burlingame, CA, USA), and incubated with primary antibodies overnight in a humidified chamber at 4C. After washing with PBS, a secondary antibody (Cat# MP-7401-15 or MP-7402, Vector Laboratories, Burlingame, CA, USA) was added for 30 min, followed by diaminobenzidine application (Cat# NC9276270, Thermo Fisher Scientific, Waltham, MA, USA) according to the manufacturer’s protocol. Samples were counterstained with hematoxylin, rinsed with distilled water, dehydrated in an increasing gradient of ethanol, cleared with xylene, and mounted with Cytoseal mounting medium (Thermo Fisher Scientific, catalog no. 8312-4). Images were recorded on a Zeiss Axioskop microscope (RRID: SCR_014587), using QCapture (RRID: SCR_014432). QuPath software (RRID: SCR_018257) was used to automatically count positive cells. For each IHC marker, five 20 × images per group were analyzed, and results were represented as the absolute number of positive cells per 20 × field. The signal was quantified by converting randomly sampled 20 × images into 16-bit images and then utilizing Fiji to employ MaxEntropy thresholding.

| Reagent | Source | Identifier | Concentration |
| --- | --- | --- | --- |
| CD4 (D7D2Z) Rabbit mAb | Cell Signaling | 25229S | 1:200 |
| CD8α (D4W2Z) XP® Rabbit mAb (Mouse Specific) | Cell Signaling | 98941 | 1:800 |
| Anti-TRAIL antibody | Abcam | ab231265 | 20 µg/ml |
| NKp46 (CD335) Polyclonal Antibody | Invitrogen | PA5-79720 | 1 µg/mL |
| FoxP3 (D6O8R) Rabbit mAb | Cell Signaling | 12653 | 1:800 |
| Granzyme B (E5V2L) Rabbit mAb | Cell Signaling | 44153 | 1:200 |
| Ki-67 (D3B5) Rabbit mAb (Mouse Preferred; IHC Formulated) | Cell Signaling | 12202 | 1:800 |
| Cleaved Caspase-3 (Asp175) Antibody | Cell Signaling | 9661 | 1:400 |
| VEGF Monoclonal Antibody (JH121) | Invitrogen | MA5-13182 | 1:20 |

### MGMT and ClpX expression analysis after imipridone with or without TMZ in DIPG cells

DIPG cells were treated at indicated doses of imipridones and/or TMZ for 48 hours in **Figure 8** before harvested for western blotting analysis. Antibodies for MGMT were obtained from Cell Signaling Technology cat #2739 and used at a titer of 1:1000 while Clpx antibody was obtained from Abcam cat #ab168338 an used at a 1:1000 dilution. β-Actin loading control antibody was obtained from Sigma-Aldrich cat #A5441 and used at a 1:5000 dilution.

### Statistical analysis

GraphPad Prism (RRID: SCR_002798) version 9.5.0 was used for statistical analyses and graphical representation (GraphPad, San Diego, CA, USA). Data are presented as means ± standard deviation (SD) or standard error of the mean (SEM). The relations between groups were compared using two-tailed, paired Student’s T-tests or one-way ANOVA tests. Survival was analyzed with the Kaplan-Meier method and was compared with the log-rank test. For Kaplan Meier curves, p-values were generated using a Survival Difference Calculator (log rank test) with default parameter ρ=0 (https://astatsa.com/LogRankTest/).

For multiple testing, Tukey’s or Benjamini-Hochberg’s methods were employed. Statistical significance is reported as follows: p ≤ 0.05: *, p ≤ 0.01: **, and p ≤ 0.001: ***.

## Acknowledgements

W.S.E-D. is an American Cancer Society Research Professor and is supported by the Mencoff Family University Professorship at Brown University. This work was supported by an NIH grant (CA173453) to W.S.E-D. This work was presented in part at the 2023 meeting of the American Association for Cancer Research.

## Supplementary Figure Legends

**Supplementary Figure 1.**
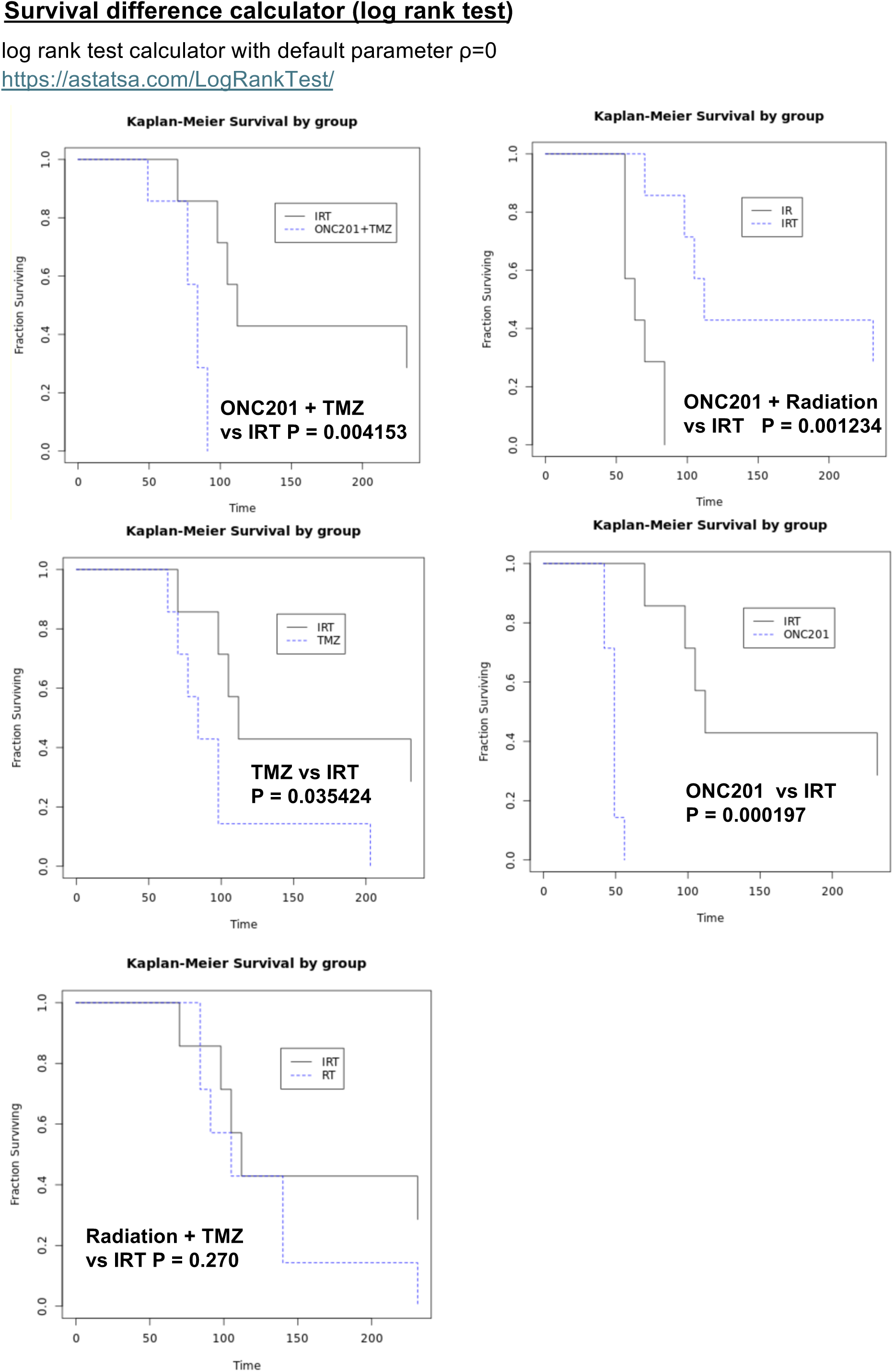
Statistical analysis of mouse survival. Survival was analyzed with the Kaplan-Meier method and was compared with the log-rank test. For Kaplan Meier curves, p-values were generated using a Survival Difference Calculator (log rank test) with default parameter ρ=0 (https://astatsa.com/LogRankTest/). Survival of treatment cohorts with mono- or doublet therapy is compared to triple IRT therapy with p-values as indicated.

